# A small molecule, ACAi-028, with anti-HIV-1 activity targets a novel hydrophobic pocket on HIV-1 capsid

**DOI:** 10.1101/2021.05.18.444758

**Authors:** Travis Chia, Tomofumi Nakamura, Masayuki Amano, Nobutoki Takamune, Masao Matsuoka, Hirotomo Nakata

## Abstract

The human immunodeficiency virus type 1 (HIV-1) capsid (CA) is an essential viral component of HIV-1 infection, and an attractive therapeutic target for antivirals. We report that a small molecule, ACAi-028, inhibits HIV-1 replication by targeting a hydrophobic pocket in the N-terminal domain of CA (CA-NTD). ACAi-028 is one of more than 40 candidate anti-HIV-1 compounds identified by in silico screening and 3-(4,5-dimethylthiazol-2-yl)-2,5-diphenyltetrazolium bromide (MTT) assays. Our binding model showed that ACAi-028 interacts with the Q13, S16, and T19 amino acid residues, via hydrogen bonds, in the targeting pocket of CA-NTD. Using recombinant fusion methods, TZM-bl, time-of-addition, and colorimetric reverse transcriptase (RT) assays, the compound was found to exert anti-HIV-1 activity in the early stage between a reverse transcriptase inhibitor, azidothymidine (AZT), and an integrase inhibitor, raltegravir (RAL), without any effect on RT activity, suggesting that this compound may affect HIV-1 core disassembly (uncoating). Moreover, electrospray ionization mass spectrometry (ESI-MS) also showed that the compound binds directly and non-covalently to the CA monomer. CA multimerization and thermal stability assays showed that ACAi-028 decreased CA multimerization and thermal stability via S16 or T19 residues.

**Importance:** These results indicate that ACAi-028 is a novel CA inhibitor that binds to the novel hydrophobic pocket of CA-NTD. This study demonstrates that a compound targeting the new hydrophobic pocket is a promising anti-HIV-1 inhibitor. The findings presented here may offer the development of a novel class of anti-viral agents that can be used, providing HIV-1 patients with more options for Anti-retroviral therapy (ART) treatment. Despite many years of successful pharmaceutical developments in the area of anti-retroviral therapy, the prevalence of drug-resistant mutations in HIV-1, necessitates the continued development of novel agents, such as ACAi-028.

## Introduction

HIV-1 is a retrovirus that has affected humans for half a century, causing 700 000 deaths and 1.7 million new infections in 2019, according to the Joint United Nations Programme on HIV and AIDS (https://www.unaids.org/en). Significant progress has been made in recent years to understand HIV infection and design drugs to counteract multiple stages of the viral life cycle. Anti-retroviral therapy (ART) has largely allowed HIV-infected patients to have the same life expectancy as uninfected individuals (1, 2). Despite of the benefits of ART, HIV-1 often acquires drug-resistant mutations, (3) resulting in treatment failure. High adherence to ART is required to sustain viral suppression during HIV clinical treatment (4). It is especially important that we continue to discover new anti-HIV-1 agents that have potent antiviral activity with different mechanisms, providing HIV patients with more options for ART treatment.

The HIV-1 capsid protein (CA) plays an essential role in both the early and late stages of the HIV-1 life cycle. One of the key periods in the early stage is HIV-1 capsid core disassembly, also known as uncoating (5, 6). After HIV-1 infection of the host cell via CD4 and CXCR4/CCR5 molecules, uncoating occurs in a regulated manner. The exact mechanism of uncoating remains unclear. Previous studies have suggested several hypotheses regarding the time and location of uncoating (7). Electron microscopy results suggest that uncoating occurs near the surface of the plasma membrane (8), while other studies have suggested that uncoating occurrs in the cytoplasm, or tethered to nuclear import (9). However, recent studies have indicated the possibility that uncoating actually occurs in the nucleus, and might be coupled with both reverse transcription and integration (10, 11, 12). A new inhibitor of CA uncoating has the potential to provide novel insights and tools for further research in this field.

Representative anti-CA compounds such as CAP-1 (13, 14), PF-3450074 (PF74) (15, 16), ebselen (17), BD-1, BM-1 (18), I-XW-053 (19), and C1 (20) have been discovered, although none have been approved for clinical use by the FDA. In this study, we used PF74 and ebselen as control drugs. PF74 is a lead compound of GS-CA1 (21) and lenacapavir, formerly known as GS-6207, (22, 23) that have been developed by Gilead Sciences, Inc. These compounds interact with the CTD-NTD interface and hinges between CA monomers, inducing hyperstabilization of the HIV-1 core and inhibiting the interaction of co-factors, such as Nup153 and CPSF6 (16). Lenacapavir has advanced to phase 2/3 of the clinical CAPELLA trial (ClinicalTrials.gov Identifier: NCT04150068), and preliminary reports from this trial indicate that it is effective at reducing the viral load of patients on failing treatment regimens, strongly suggesting that the CA is a viable target for the development of novel anti-HIV-1 compounds. Ebselen is also a unique CA inhibitor that covalently binds to the C-terminal domain of CA via Cysteine residues, inhibiting HIV-1 activity (17).

We previously reported that the insertion of a short amino acid sequence near the Arg18/Thr19 region of decreased its stability, inducing abnormal CA degradation (24), and identified an adequate hydrophobic cavity near this region on the surface of CA-NTD, which could be a potential drug target. To the best of our knowledge, none of the other published anti-CA compounds are known to target this hydrophobic cavity.

In this study, we searched for new compounds capable of interacting with the hydrophobic pocket from a library containing millions of commercially available compounds via *in silico* docking simulations. We identified several compounds as candidate HIV-1 inhibitors that were capable of preventing HIV-1 replication. Here, we highlight ACAi-028, a CA inhibitor with unique molecular characteristics, which possess potent anti-HIV-1 activity. It is anticipated that both this compound and the identified binding pocket hold potential therapeutic and research applications.

## Results

### Identification of candidate compounds that target a novel hydrophobic cavity of the HIV-1 capsid

CA protein consists of the N-terminal domain (CA-NTD, amino acid residues 1-145) and the C-terminal domain (CA-CTD, residues 151-231) linked via a short flexible region (residues 149-150) (25, 26, 27). CA-NTD comprises one β-hairpin, seven α-helices, and a cyclophilin binding loop (CypA-BL), while CA-CTD has a 3_10_-helix and four α-helices, as shown in Fig. 1A. Recently, CA inhibitors have attracted attention due to the development of lenacapavir, whose lead compound is PF74 (15), which exhibits a long-acting, strong anti-HIV profile. We have reported that the insertion of a short amino acid sequence into the CA-NTD, specifically near the R18 and T19 residues, results in spontaneous CA degradation (24).

**Figure 1.**
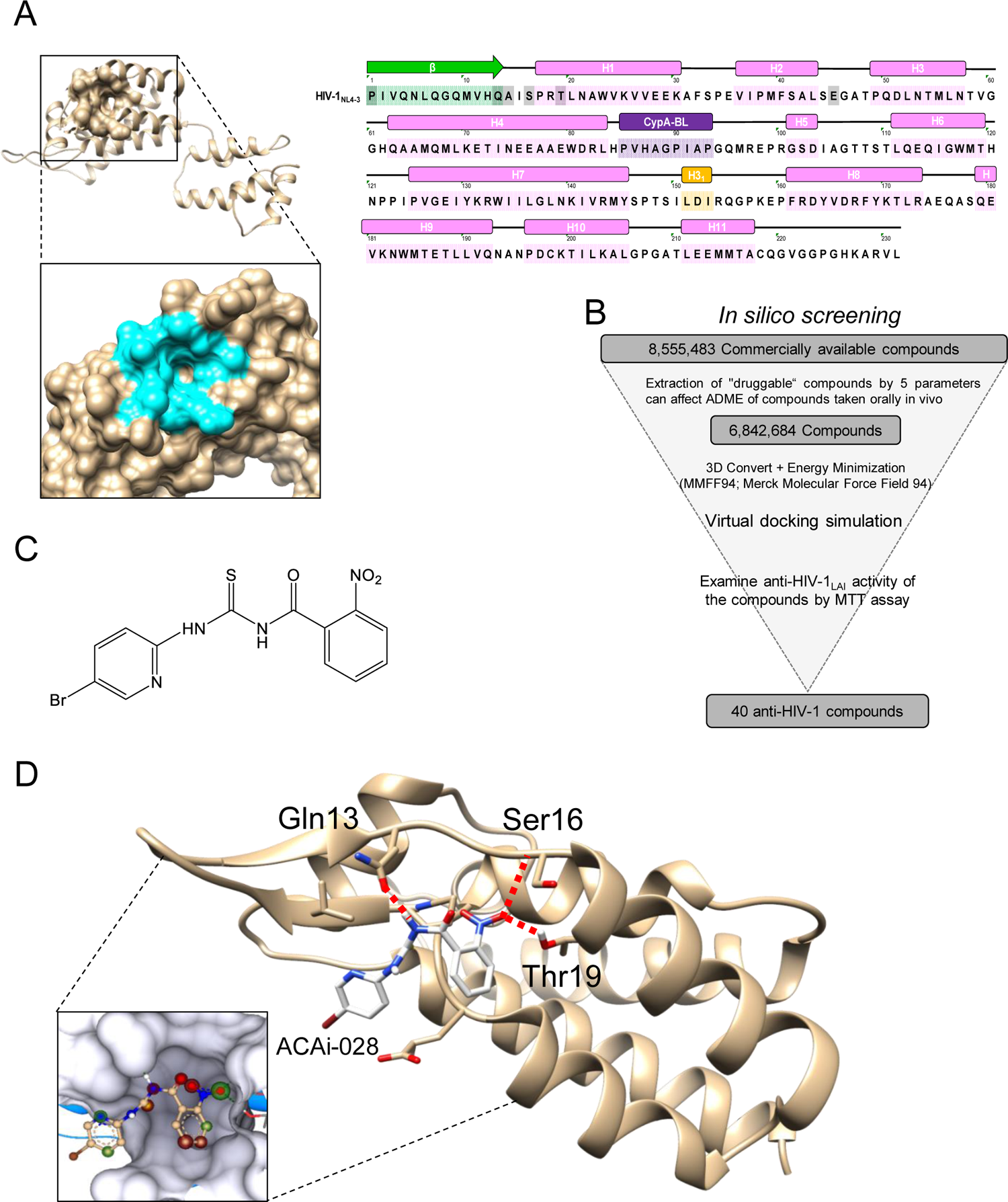
Profile of CA, target cavity, and *in silico* docking simulations of ACAi-028. (**A)** 3D structures of full-length CA (PDB accession number 4XFX) shown in tan, and closed-up of ACAi-028-targeted cavity in the CA-NTD shown in light blue are on the left side. The CA comprises a β-hairpin in green, CypA-BL in purple, eleven α-helices: H1-11 in pink, and a 3_10_-helix in yellow on the right side. (**B**) The procedure of *in silico* screening to identify as candidate anti-CA inhibitors. (**C**) The chemical structure of ACAi-028. (**D**) The docking simulation result of ACAi-028 with the target cavity is shown. Hydrogen bond interactions between molecular surface of CA CA-NTD _1-146/Δ87-99G_ crystal and ACAi-028 are indicated in dotted lines. The carbons of CA and ACAi-028 are shown in tan and white colors, respectively. Nitrogen atoms, oxygen atoms, hydrogen atoms, and bromine atoms are shown in blue, red, white, and brown, respectively. ACAi-028 forms two H-bonds interaction with the side-chains of Gln13 and Thr19 (inter-atomic distances of 1.87 Å and 1.89 Å, respectively) and H-bond interaction with the main-chain of Ser16 (inter-atomic distance of 2.06 Å). Molecular graphics was performed with UCSF Chimera (https://www.rbvi.ucsf.edu/chimera). All docking simulations were performed with SeeSAR and FlexX version 10 (BioSolveIT GmbH, Sankt Augustin, Germany).

To identify new capsid inhibitors, we selected a novel hydrophobic pocket (Fig. 1A) near this position as a potential interaction site for drug candidates with the CA-NTD. *In silico* docking simulations were performed to search for new compounds that interact with this pocket from a database of over eight million commercially available compounds. The selection process is illustrated in Figure 1B. More than 40 compounds were identified as candidate HIV-1 inhibitors that prevented HIV-1_LAI_ replication, using the MTT assay.

In this study, we identified ACAi-028 (Fig. 1C), which has a small molecular weight (MW) of 381 g/mol, and potent anti-HIV-1_LAI_ activity (EC_50_, 0.55 µM), among the candidate HIV-1 inhibitors. Therefore, we examined the anti-HIV profile of ACAi-028.

The crystal structure of CA-NTD_1-146/Δ87-99G_ was produced (15) (Fig. S1), and the binding profile of ACAi-028 to the CA-NTD was elucidated using a docking model (Fig. 1D). The binding profile of ACAi-028 with the CA-NTD is similar to that of ACAi-028 with full-length CA (Protein Data Bank [PDB] accession number 4XFX) (28) (Fig. S2). ACAi-028 is estimated to form two hydrogen bonding (H-bond) interactions with the side-chains of Q13 and T19 (inter-atomic distances of 1.87 and 1.89 Å, respectively), and one H-bond interaction with the main chain of S16 (inter-atomic distance of 2.06 Å). This structural arrangement suggests that ACAi-028 could potentially bind to the newly identified hydrophobic cavity in CA-NTD.

### ACAi-028 inhibits the early stage of the HIV-1 life cycle

Next, we determined the anti-HIV activity (EC_50_s) of ACAi-028 against various HIV-1 and HIV-2 strains, in comparison with several compounds that have been previously reported as capsid inhibitors, such as PF74 (15) and ebselen (17), and an RT inhibitor, AZT (29) (Table 1).

**Table 1:** Comparison of anti-HIV-1 activity of ACAi-028 against other inhibitors.

| Drugs | EC<sub>50</sub> (μM) |  |  |  |
| --- | --- | --- | --- | --- |
|  | Single round inhibition<sup>a</sup> | Multiple round inhibition<sup>b</sup> |  |  |
|  | TZM-bl | MT-4 | MT-2 |  |
|  | HIV-1<sub>VSVG-NL4-3</sub> | HIV-1<sub>NL4-3</sub> | HIV-1<sub>LAI</sub> | HIV-2<sub>ROD</sub> |
| ACAi-028 | 0.48 ± 0.21 | 0.12 ± 0.04 | 0.55 ± 0.04 | >10 |
| PF74 | 0.76 ± 0.55 | 0.33 ± 0.08 | 0.37 ± 0.50 | >10 |
| Ebselen | 1.68 ± 1.45 | 1.83 ± 0.10 | 1.73 ± 0.04 | n.d. |
| AZT | 0.14 ± 0.13 | 0.26 ± 0.02 | 0.23 ± 0.05 | n.d. |

<sup>a</sup> TZM-bl cells (5×10<sup>4</sup>/mL) were subjected to HIV-1<sub>VSVG-NL4-3</sub> (50 ng/mL of p24) in 96-well plates for 48 h in the presence of the tested compounds. The supernatant was removed and lysis buffer added to the samples. After the samples were shaken for 25 minutes, D-luciferin was added to each well. The luciferase intensity was measured using FluoSTAR Omega.ssay at 48 h. <sup>b</sup> MT-2 cells (10<sup>4</sup>/ml) were exposed to 100 TCID<sub>50</sub> of HIV-1<sub>LAI</sub> or HIV-2<sub>ROD</sub> and cultured in the presence of various concentrations of each compound, and the EC<sub>50</sub> (50% effective concentration) values were determined by the MTT assay. For HIV-1<sub>NL4-3</sub>, the EC<sub>50</sub> values were determined by using MT-4 cells as target cells. MT-4 cells (10<sup>5</sup>/ml) were exposed to 100 TCID<sub>50</sub>s of HIV-1<sub>NL4-3</sub>, and the inhibition of p24 Gag protein production by each drug was used as an endpoint. All assays were conducted in duplicate, and the data shown represent mean values (± 1 S.D.) derived from the results of two or three independent experiments.

ACAi-028 exerted potent anti-HIV-1 activity against single-round HIV-1 infection using VSV-G pseudo-typed HIV-1_NL4–3_ (HIV-1_VSV-G dENV_), suggesting that ACAi-028 inhibits the early stage of the HIV-1 life cycle. ACAi-028 also prevented multiple-round HIV-1 infection using HIV-1 strains (HIV-1_NL4-3_ and HIV-1_LAI_), except for HIV-2 strain (HIV-2_ROD_), at a concentration similar to that of PF74, ebselen, and AZT. In addition, ACAi-028 inhibited the replication of various types of HIV-1 strains (HIV-1_104pre_, HIV-1_MDR/B_ (30) in peripheral blood mononuclear cells (PBMCs) and HIV-1_ATV_^R^_5µM_ in MT-4 cells (Table 2), and had negligible cytotoxicity in all the cell lines we tested (Table 3).

**Table 2:**
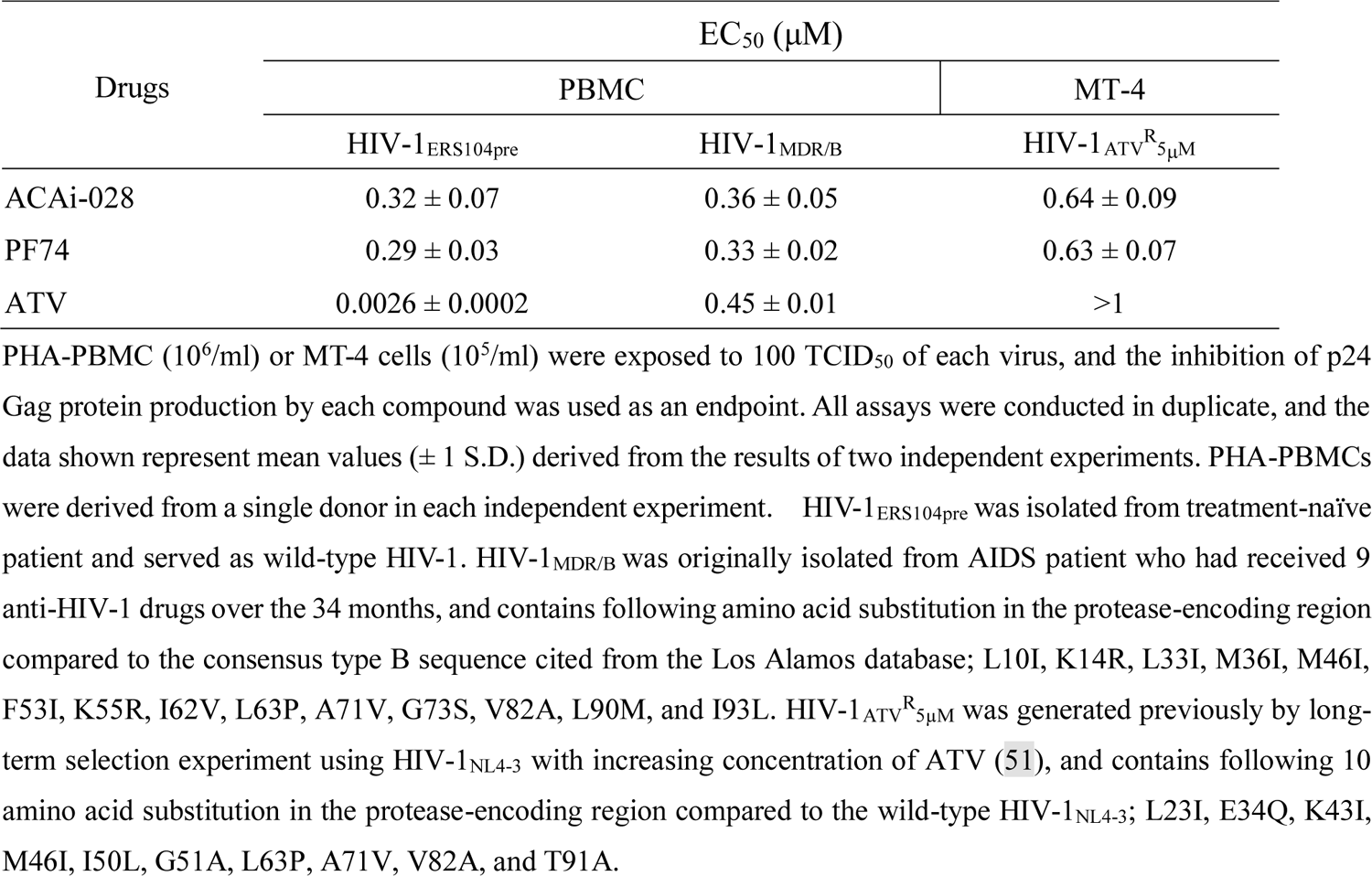
Anti-HIV-1 activity of ACAi-028 against clinically isolated multi-drug-resistant HIV-1 in PBMC and protease-inhibitor-resistant laboratory strain in MT-4 cells.

| Drugs | EC<sub>50</sub> (μM) |  |  |
| --- | --- | --- | --- |
|  | PBMC |  | MT-4 |
|  | HIV-1<sub>ERS104pre</sub> | HIV-1<sub>MDR/B</sub> | HIV-1<sub>ATV<sup>R</sup>5μM</sub> |
| ACAi-028 | 0.32 ± 0.07 | 0.36 ± 0.05 | 0.64 ± 0.09 |
| PF74 | 0.29 ± 0.03 | 0.33 ± 0.02 | 0.63 ± 0.07 |
| ATV | 0.0026 ± 0.0002 | 0.45 ± 0.01 | >1 |

PHA-PBMC (10<sup>6</sup>/ml) or MT-4 cells (10<sup>5</sup>/ml) were exposed to 100 TCID<sub>50</sub> of each virus, and the inhibition of p24 Gag protein production by each compound was used as an endpoint. All assays were conducted in duplicate, and the data shown represent mean values (± 1 S.D.) derived from the results of two independent experiments. PHA-PBMCs were derived from a single donor in each independent experiment. HIV-1<sub>ERS104pre</sub> was isolated from treatment-naïve patient and served as wild-type HIV-1. HIV-1<sub>MDR/B</sub> was originally isolated from AIDS patient who had received 9 anti-HIV-1 drugs over the 34 months, and contains following amino acid substitution in the protease-encoding region compared to the consensus type B sequence cited from the Los Alamos database; L10I, K14R, L33I, M36I, M46I, F53I, K55R, I62V, L63P, A71V, G73S, V82A, L90M, and I93L. HIV-1<sub>ATV<sup>R</sup>5μM</sub> was generated previously by long-term selection experiment using HIV-1<sub>NL4-3</sub> with increasing concentration of ATV (51), and contains following 10 amino acid substitution in the protease-encoding region compared to the wild-type HIV-1<sub>NL4-3</sub>; L23I, E34Q, K43I, M46I, I50L, G51A, L63P, A71V, V82A, and T91A.

**Table 3:** Comparison of ACAi-028 cytotoxicity against other inhibitors in various cell lines.

| Cells | CC<sub>50</sub> (μM) |  |  |
| --- | --- | --- | --- |
|  | ACAi-028 | PF74 | Ebselen |
| Li-7 (hepatocyte) | >100 | >100 | >20 |
| HLE (hepatocyte) | 72.3 | >100 | >20 |
| HEK293 (kidney) | 62.6 | >100 | >20 |
| MT-4 (lymphocyte) | >100 | >100 | >20 |
| MT-2 (lymphocyte) | 86.6 | >100 | >20 |
| PBMC | >100 | n.d. | n.d. |

Cells were incubated with the respective compounds for one week before cytotoxicity was quantified using MTT assay.

To examine the details of early stage inhibition by ACAi-028, we performed fusion (31), TZM-bl, time-of-addition (32), and colorimetric RT activity assays. ACAi-028 did not significantly inhibit the entry step of HIV-1_LAI_, in comparison to the entry inhibitor, AMD3100 (33) (Fig. 2A). ACAi-028 also displayed strong anti-HIV-1 activity against HIV-1 R5 strains, including HIV-1_Ba-L_, and HIV-1_JR-FL_, and X4 strains, except for HIV-2_ROD_, using the TZM-bl assay (Fig. 2B).

**Figure 2.**
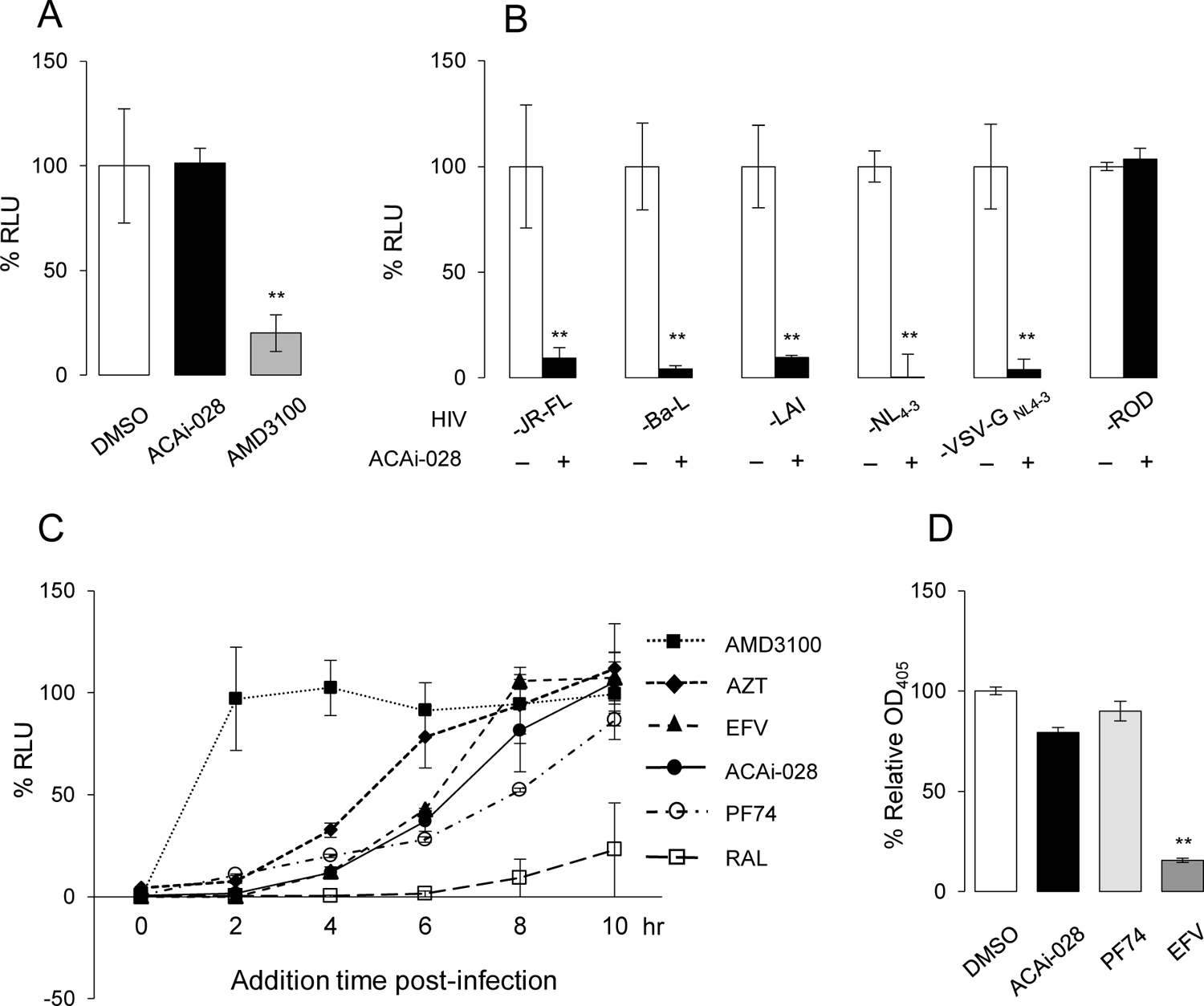
The early-stage inhibition of ACAi-028 in the HIV-1 life cycle. **(A)** Fusion assay was conducted between 293T and COS-7 cells in the presence of ACAi-028 or AMD3100. Results of the luciferase intensity are shown as percentage compared to Dimethyl sulfoxide (DMSO) controls. AMD3100 is a fusion inhibitor against CXCR4 co-receptors of X4-tropic HIV-1 (B) The early-stage inhibition of ACAi-028 against X4-tropic (HIV-1_LAI_ and HIV-1_NL4-3_), R5-tropic (HIV-1_Ba-L_ and HIV-1_JR-FL_), VSV-G HIV-1_NL4-3_, and HIV-2_ROD_ strains were measured using TZM-bl cells. Luciferase activity was measured at 48 h post-infection and is shown as a percentage normalized to DMSO controls. (**C**) Time-of-addition assay was conducted by the addition of ACAi-028 and various early-stage inhibitors such as AMD3100, AZT, EFV, PF74, and RAL to TZM-bl cells in 2 h intervals up to 10 h. Data was shown as percentage every 2 h and normalized to DMSO controls (**D**) Effects of ACAi-028, PF-74, and EFV on RT activity were measured by colorimetric reverse transcriptase assay. OD_405_ values were measured and shown as percentages. All assays were performed in duplicate, and error bars indicate ±SD from at least two independent experiments. Statistical significance was examined using Student’s t-test; *, P < 0.05, **, P < 0.005.

A time-of-addition assay was conducted to compare the inhibition time of ACAi-028 with that of the other classes of anti-HIV-1 drugs. As expected, AMD3100 ceased displaying anti-HIV-1 activity when it was added more than 2 h post-infection, while an integrase inhibitor, RAL, significantly prevented HIV-1 infection beyond10 h post-infection (Fig. 2C).

We observed that ACAi-028 inhibited anti-HIV-1 activity at a similar time as RT inhibitors, AZT and efavirenz (EFV) (34), as well as a CA inhibitor, PF74. Fifty percent inhibition induced by these drugs occurred between 4 and 8 h post-infection, which is consistent with a previous report (15). Moreover, we examined whether ACAi-028 prevents HIV-1 RT activity using a colorimetric RT activity assay and found that ACAi-028 and PF74 did not have a significant impact on HIV-1 RT activity, in comparison to EFV (Fig. 2D) *in vitro*. This suggests that ACAi-028 is unlikely to be an RT inhibitor. Taken together these results indicate that ACAi-028 may have a similar level of potency as a CA inhibitor as PF74, which affects the CA uncoating process.

### ACAi-028 does not affect the late stage of the HIV-1 life cycle

To examine the effect of ACAi-028 on late-stage inhibition, we examined whether ACAi-028 affects the process of HIV-1 production, including Gag proteolytic processing and maturation. Forty-eight hours after transfection of pHIV-1_NL4-3_ into 293T cells in the presence of ACAi-028 (10 µM), PF74 (10 µM), ebselen (10 µM) or, a protease inhibitor, darunavir (DRV) (2 µM) (35). Gag proteins within the cells were observed by western blotting, and viral production was evaluated comparison to p24 expression levels in the culture supernatant. ACAi-028, PF74, and ebselen did not affect Gag proteins in the cells, unlike DRV (Fig. 3A), whereas ACAi-028 (88.7%), and ebselen (75.2%), PF74 (62.9%), and DRV (15.7%) reduced HIV-1 production, which is consistent with previous reports (15, 17) (Fig. 3B). Additionally, the effect of ACAi-028 on Gag proteolytic processing and maturation of HIV-1 virions was investigated using western blotting and the TZM-bl assay. ACAi-028 did not affect HIV-1 maturation, as observed via western blotting, using anti-Gag or anti-IN antibodies as well as a DMSO control (Fig. 3C). ACAi-028 did not reduce the viable virions (111.8%), as opposed to DRV (3.6%) (Fig. 3D), suggesting that ACAi-028 does not have any effect on HIV-1 production and maturation. On the other hand, a high concentration of PF74 (10 µM) did not affect Gag proteins in the cell lysate (Fig. 3A), but the production level of virions in the presence of PF74 significantly decreased by 62.9% compared to that of the DMSO control. Furthermore, the infectivity of the virions was reduced to 6.9% (Fig. 3B and D), suggesting that PF74 affects both the early and late stages, in agreement with previous reports (15). It has been reported that ebselen does not exhibit late-stage inhibition (17). Interestingly, premature products such as Gag-pol (p160), Gag-pol intermediate (p120), Gag (p55), and Gag intermediates were found in the virion lysate that was produced at a high concentration (10 µM) of Ebselen, via the western blotting with anti-Gag or anti-IN antibodies, which is similar to DRV (Fig. 3C). Additionally, the infectivity of the virions produced in the presence of ebselen (10 µM) was significantly reduced by 64.6 % of their normal level (Fig. 3D). These results suggest that ebselen may interfere with HIV-1 maturation. These results indicate that ACAi-028 is unlikely to affect the late stage of the HIV-1 life cycle.

**Figure 3.**
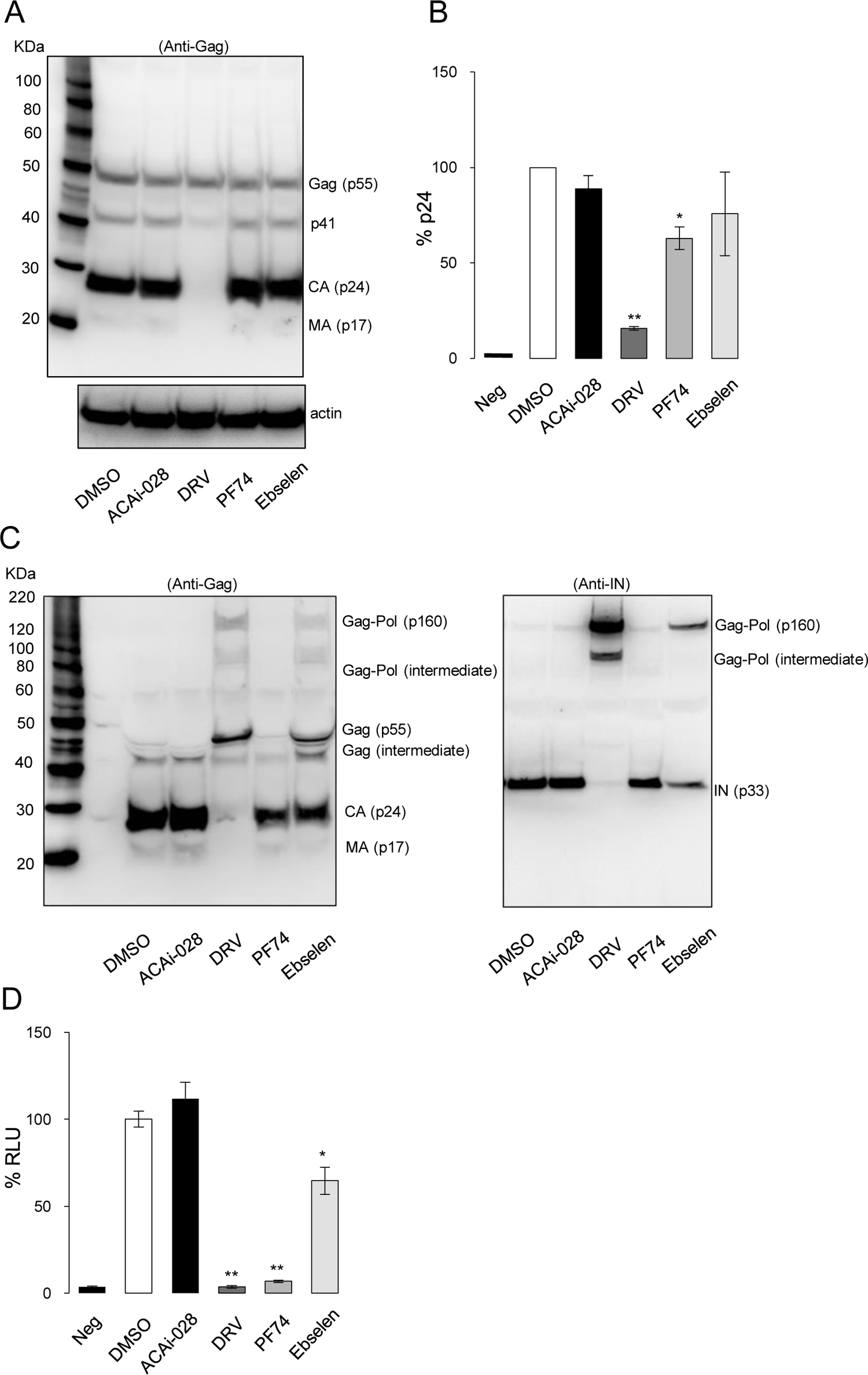
Effect of ACAi-028 on the latestage of the HIV-1 life cycle. (**A**) Gag-pol proteolytic processing and virus production were examined by the western blotting with anti-Gag antibody in the cell lysate and (**B**) measuring the p24 levels in the supernatant of 293T cells which were transfected with pNL_4-3_ in the presence of ACAi-028 (10 µM), PF74 (10 µM), Ebselen (10 µM), or DRV (2 µM). After the virions in the supernatants were purified, Gag-pol proteolytic processing and HIV-1 maturation of the virions were examined by (**C**) the western blotting with anti-Gag on the left side and anti-IN antibody on the right side, and by (**D**) TZM-bl assay at the bottom. All assays were performed in duplicate, and error bars indicate ±SD from three independent experiments. Statistical significance was examined using Student’s t-test; *, P < 0.05, **, P < 0.005.

### Conformational difference of targeting the hydrophobic pocket between HIV-1 and HIV-2

ACAi-028 did not show anti-HIV activity against the HIV-2_ROD_ strain (Table 1). We compared the amino acid sequences of experimental HIV-1 strains, such as HIV-1_NL4-3_, HIV-1_HXB (LAI)_, HIV-1_Ba-L_, and HIV-1_JR-FL_, which are all closely related to simian immunodeficiency virus from chimpanzees (SIV cpz). Similarly, comparisons of experimental HIV-2 strains, such as HIV-2_ROD_ and HIV-2_EHO_, which are related to SIV sooty mangabeys (SIV smn), as shown in Fig. 4A, were also undertaken. The Q13, S16, and T19 residues, which are expected to play important roles in ACAi-028 binding to CA (Fig. 1D), are conserved among all HIV-1 strains and SIV cpz, whereas most of the amino acids from residues 2 to 15 of HIV-2 are different from those of HIV-1 (Fig. 4A). Moreover, Q13, S16, and T19 residues were highly conserved across 6144 sequences from all HIV-1 subtypes (HIV sequence database filtered web alignment) at conservation rates of 99.89%, 97.85%, and 99.72 %, respectively (Fig. 4B).

**Figure 4.**
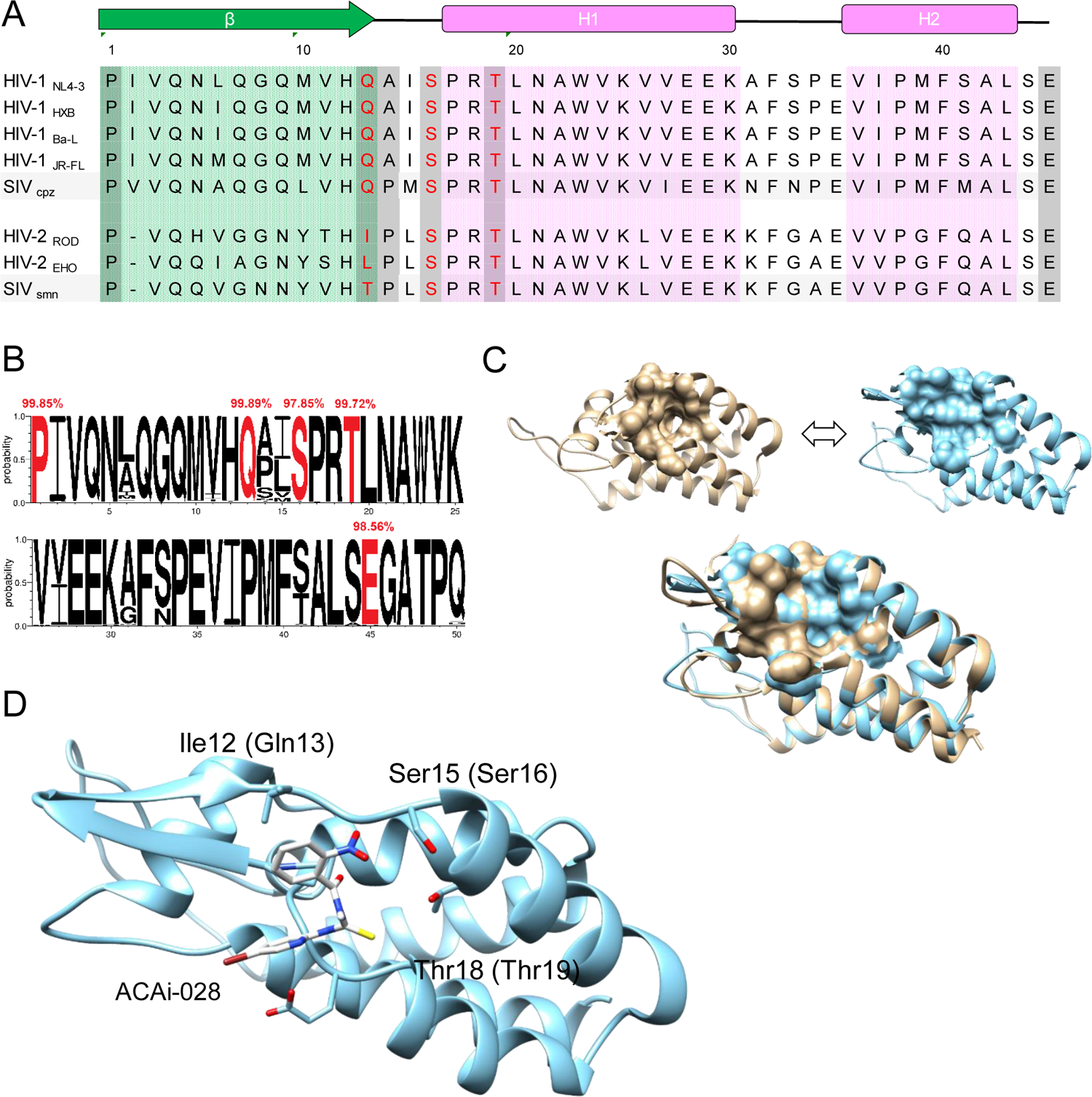
Amino acid residues and structures of CA-NTD among HIV and SIV. **(A)** Alignment of CA-NTD (residues from 1 to 45) among laboratory HIV-1 strains, SIV_CPZ_, HIV-2 strains, and SIV_smn_. (**B**) Representation of frequencies of CA-NTD (residues from 1 to 50) from 6144 sequences of all HIV-1 subtypes (Los Alamos HIV sequence database filtered web alignment) using the WebLogo 3.7.4 application (http://weblogo.threeplusone.com/create.cgi). Highlighted in red are the five residues of (Pro1, Gin13, Ser16, Thr19, and Glu45) that constitute the ACAi-028 binding cavity. Above each of these residues is the percentage consensus as determined by Jalview 2.11.1.4 program (http://www.jalview.org) (52). (**C**) The target cavity on the surface of CA-NTD_1-146/Δ87-99G_ monomer used in the flexible docking simulation is shown in tan in the left panel. CA-NTD_HIV-2_ (CA-NTD_ROD_) monomer ([PDB] accession number 2WLV) and corresponding target cavity is shown in cyan in the right panel. The bottom panel shows merged image of the CA-NTD of both strains. (**D**) The docking simulation result of ACAi-028 with the target cavity in CA-NTD_ROD_ is shown. Hydrogen bond interactions between molecular surface of CA-NTD_ROD_ and ACAi-028 were not observed unlike the docking result between CA-NTD_HIV-1_ and ACAi-028. The carbons of CA and ACAi-028 are shown in cyan and white colors, respectively. Nitrogen atoms, oxygen atoms, hydrogen atoms, and bromine atoms are shown in blue, red, white, and brown, respectively. Docking simulations were performed with SeeSAR and FlexX version 10. Molecular graphics was performed with UCSF Chimera.

The crystal structures of the ACAi-028 target hydrophobic pocket of HIV-1_NL4-3_ ([PDB] accession number 4XFX) (28) and HIV-2_ROD_ ([PDB] accession number 2WLV) (36) are shown in Fig. 4C. This HIV-1_NL4-3_ pocket seems to have a sufficient volume for ACAi-028 binding, while that of HIV-2_ROD_ appears to be too shallow for binding (Fig. 4C and Fig. S3). The cavity of CA-NTD_NL4-3_ is covered by that of the CA-NTD_ROD_ in the overlay of these crystal structures, as shown in Fig. 4C, suggesting that the ACAi-028 target volume of HIV-2 is clearly smaller than that of HIV-1.

Moreover, we examined the binding ability of ACAi-028 to CA-NTD_ROD_ using a binding model. As shown in Fig. 4D, there were no bridging H-bonds between ACAi-028 and the amino acid residues of CA-NTD _ROD_, corresponding to the lack of inhibition (EC_50_ > 10 µM) of ACAi-028 against HIV-2 _ROD_ (Table 1). These results indicate that ACAi-028 may fail to interact with HIV-2_ROD_ CA, resulting in no anti-HIV-2 activity.

### Binding profiles of ACAi-028 to CA proteins

In order to observe the direct binding of ACAi-028 to CA, we produced recombinant HIV-1_NL4-3_-derived CA proteins using *E. coli*, and examined the binding of ACAi-028 to these proteins using an electrospray ionization mass spectrometry (ESI-MS) (37). The ESI-MS spectra of the CA monomer with 1% methanol revealed nine peaks of charged ions in the range of mass/charge ratio (*m/z*) range of 1,100–1,900 (Fig. 5A). The MW estimated from the peaks of charged irons (deconvoluted ESI-MS spectrum) was 25601.9 Da, corresponding to the theoretical MW of intact CA monomer (25602.5 Da), as calculated by Peptide Mass Calculator v3.2 (Fig. 5A). After treatment of CA with ACAi-028, peaks associated with ACAi-028 binding to CA emerged in the *m/z* range of 1,300–1,900 next to each spectrum of the CA monomer (Fig. 5B). The deconvoluted ESI-MS spectrum revealed a peak associated with CA and ACAi-028 at 25984.1 Da, which was similar to the sum of the MW of a CA monomer (25601.9 Da) and ACAi-028 (381 Da) (Fig. 5B). Additionally, we examined whether ACAi-028 binds covalently to CA. After treatment of CA with ACAi-028 or ebselen, the samples were denatured by acetonitrile and trifluoroacetic acid. The binding peak of CA with ACAi-028 was not detected in Fig. 5C, while two strong peaks of CA with ebselen (25922.03 and 26120.59 Da, representing CA combined with one or two ebselen molecules, respectively) were observed (Fig. 5D).

**Figure 5.**
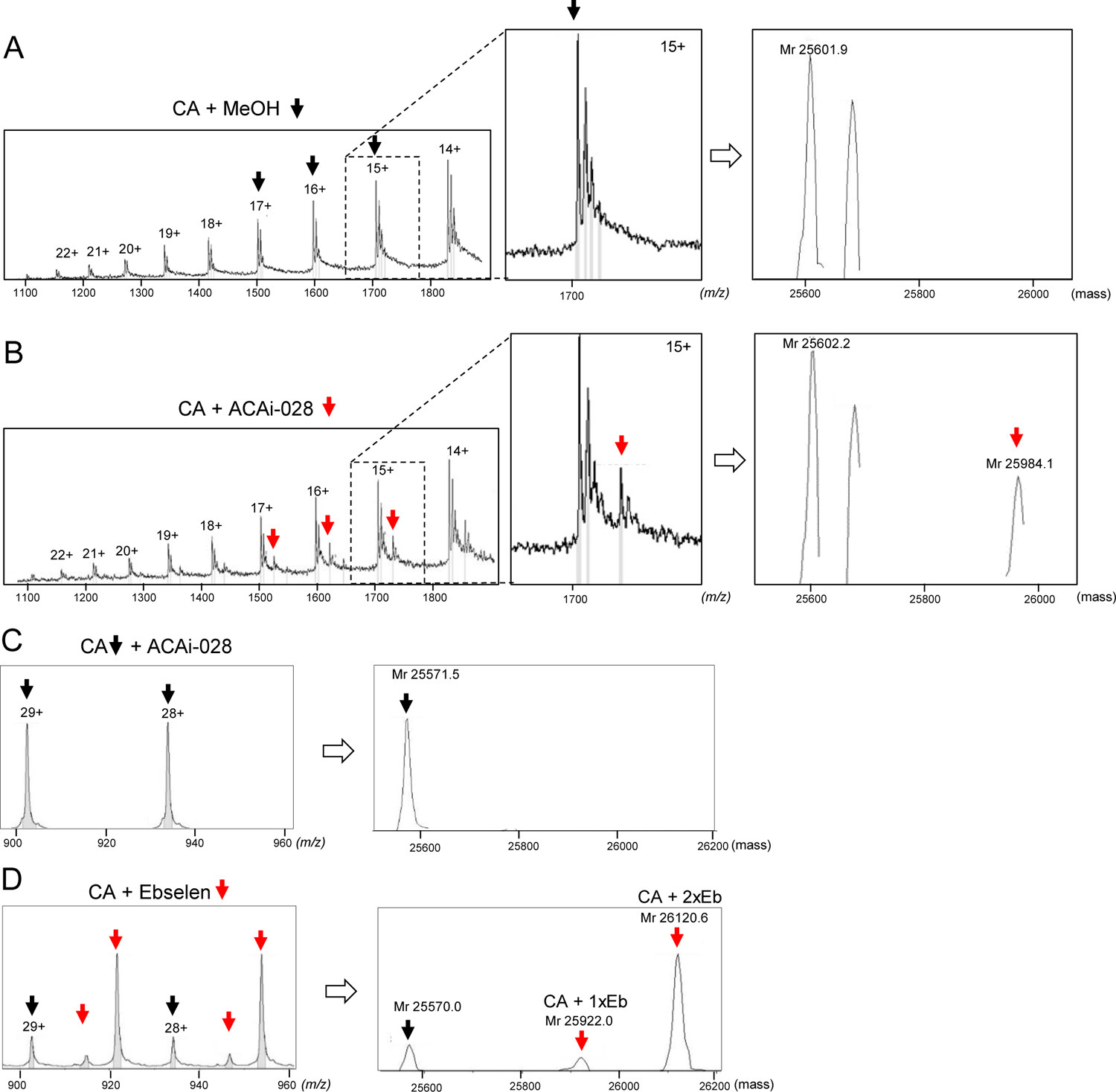
ACAi-028 interacts directly and non-covalently with CA monomer ESI-MS spectra of ACAi-028 (50 µM) and Ebselen (50 µM) binding to CA. (**A**) Black arrows represent CA monomer peaks of the charged ions. Deconvoluted ESI-MS spectrum of CA monomer is shown on the right side. (**B**) ESI-MS spectra of CA binding to ACAi-028 are shown in red arrows. Deconvoluted ESI-MS spectra of CA monomer or CA monomer binding to ACAi-028 are shown on right side. (**C**) Under denaturing condition, acetonitrile and trifluoroacetic acid are added to a mixture CA of ACAi-028. Black arrows indicate CA monomers failed to interact with ACAi-028. Deconvoluted ESI-MS spectra is shown on the right side. (**D**) Under the same condition as (**C**), black arrows indicate CA monomers and red arrows represent CA monomer binding covalently to one (left) or two (right) ebselen, respectively. Each deconvoluted ESI-MS spectrum is shown on the right side.

These results are in line with a previous report (17), suggesting that ACAi-028 binds non-covalently to CA, unlike ebselen. Thus, ACAi-028 binds directly and non-covalently to the CA monomer.

### ACAi-028 affects the molecular characterization of CA proteins via S16 and T19 residues

ACAi-028 was shown to interact directly with the CA proteins using ESI-MS. To investigate how ACAi-028 interacts with CA proteins, we produced CA variants carrying S16E (CA_S16E_) or T19A (CA_T19A_) amino acid substitutions, which were intend to alter the binding ability of ACAi-028 to CA.

CA_M185A_, carrying an M185A amino acid substitution was also produced as previously reported (38), as a control. CA thermal stability in the presence of ACAi-028 was examined using differential scanning fluorimetry (DSF) (39, 40) (Fig. 6). The melting temperature (Tm) value for proteins generally indicates the temperature at which the protein is unfolded by 50% folded (Tm 50). Tm 50 of wild-type CA (CA_WT_) increased in the presence of PF74, while that of ACAi-028 clearly decreased by 6.8°C and 7.1°C at 10 and 50 µM, respectively (Fig. 6A and B). Tm 50 of CA_S16E_ showed a mild reduction of 3.9°C at 50 µM, while that of CA_T19A_ was seen to remain unchanged in the presence of ACAi-028 (Fig. 6C and D), suggesting that S16 and T19 residues may be associated with the binding of ACAi-028 to the target pocket of CA-NTD. Additionally, a CA multimerization assay was performed in the presence of ACAi-028. The treatment of CA_WT_ with PF74 increased CA multimerization as previously described (15), whereas ACAi-028 decreased CA_WT_ multimerization at 4 and 40 µM in a concentration-dependent manner (Fig. 7A). M185A greatly decreased the CA multimerization, in agreement with, a previous report (38). The addition of S16E substitution to CA proteins slightly reduced CA_S16E_ multimerization, while T19A slightlyincreased CA_T19A_ multimerization in comparison to CA_WT_ (Fig. 7B) at the same sodium concentration, suggesting that these residues significantly affect the CA multimerization. In the presence of ACAi-028, CA_S16E_ multimerization was similar to that of CAWT (Fig. 7C). However, CA_T19A_ multimerization was not affected by the presence of ACAi-028, even at a higher concentration (40 µM) (Fig. 7D), suggesting that the T19A mutant may inhibit ACAi-028 binding to the target pocket under these conditions. These results strongly indicate that ACAi-028 may interact with CA proteins via S16 and T19 residues in the target pocket of CA-NTD.

**Figure 6.**
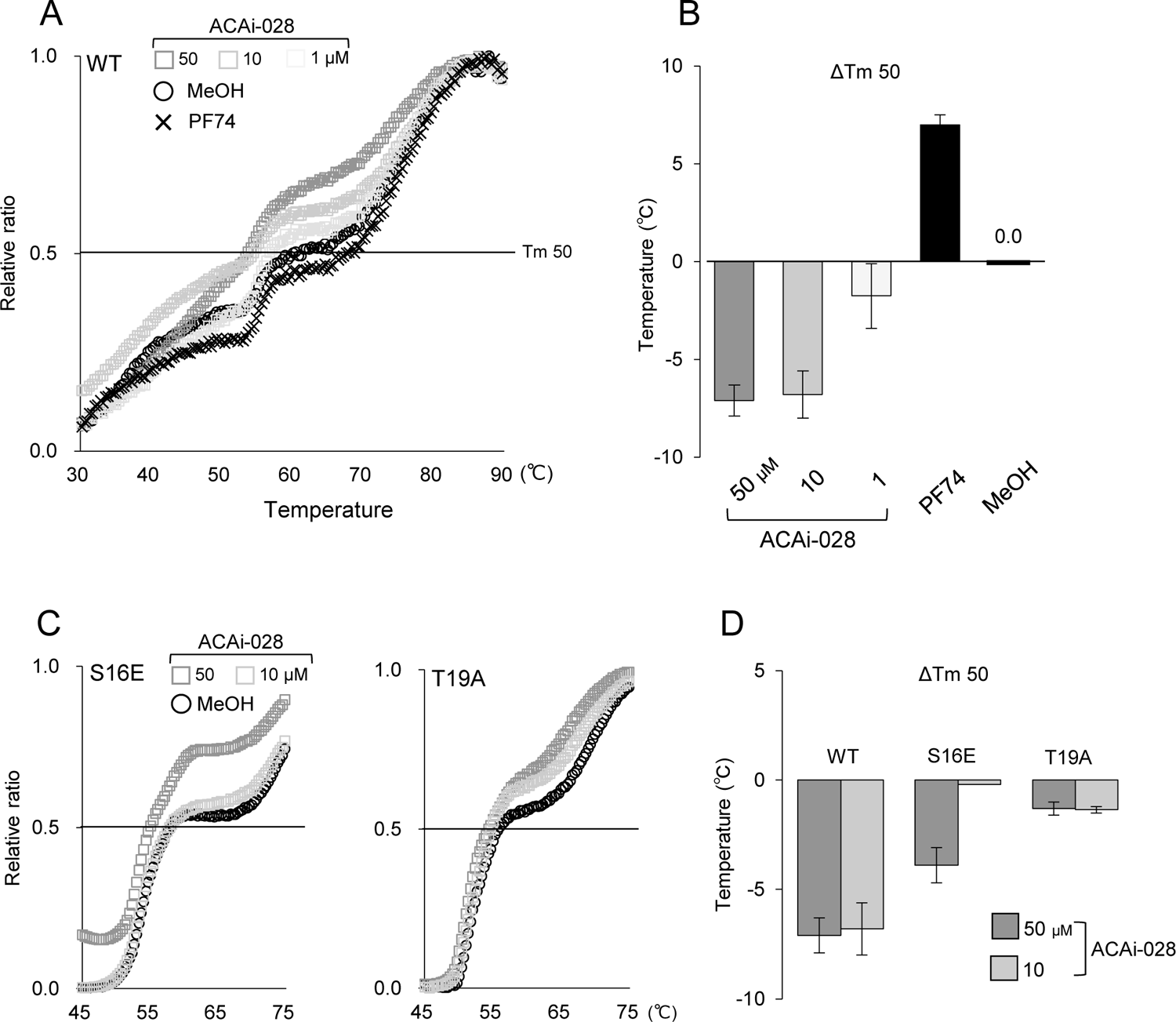
Effect of ACAi-028 on CA thermal stability. (**A**) DSF was performed using SYBR-Orange dye in the presence of methanol shown in black circle, a different concentration (1, 10, and 50 µM) of ACAi-028 shown in grey squares, and PF74 (10 µM) shown in black triangle to examine CA stability. (**B**) Graphical representation of Tm changes (ΔTm 50) from (**A**) are shown for ACAi-028 and PF74. (**C**) DSF of CA_S16E_ and CA_T19A_ were tested in high concentration of ACAi-028 (10 µM and 50 µM shown in light and dark grey squares, respectively). Black circle indicates in methanol as a control. (**D**) Graphical representation of ΔTm 50 from (**C**) are shown for (10 and 50 µM) ACAi-028. All assays were performed in triplicate, and error bars indicate ±SD from three independent experiments. Statistical significance was examined using Student’s t-test (*, P < 0.05, **, P < 0.005).

**Figure 7.**
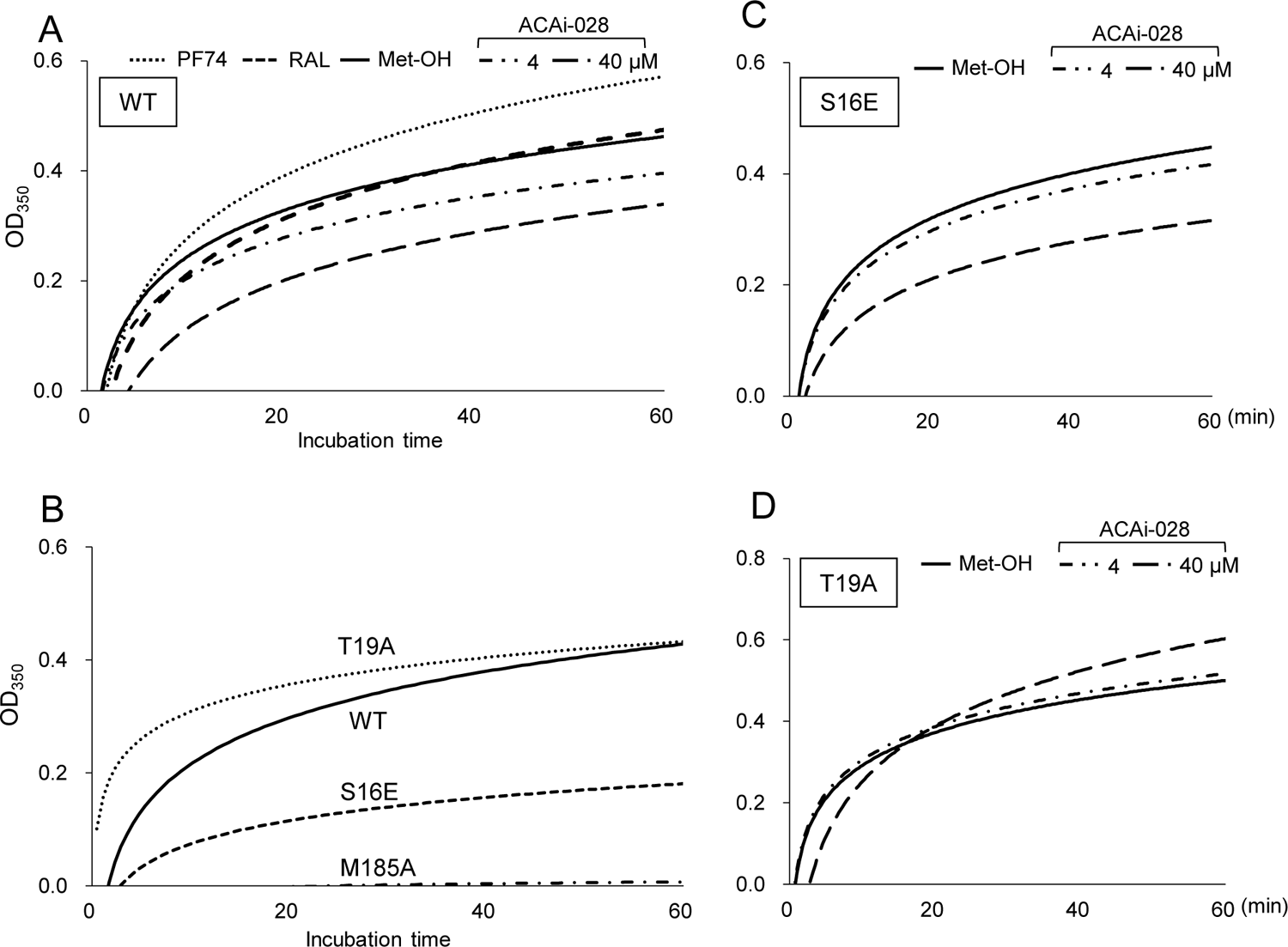
Effect of ACAi-028 on CA multimerization. (A) CA multimerization assay was performed by the addition of a high sodium buffer (the ratio of 150 mM sodium phosphate to 5M sodium phosphate buffer is five to five) in the presence of ACAi-028 (4 and 40 µM), PF74 (4 µM), and RAL (40 µM). Turbidity of the mixtures was measured at OD_350_ over a period of 60 minutes. PF74 and RAL are used as positive or negative controls, respectively. Representative data is shown from three independent experiments. (**B**) CA _WT_ multimerization was compared to CA_T19A_, CA_S16E_, and CA_M185A_ multimerization (The ratio of 150 mM to 5M sodium is five to five). Effects of ACAi-028 (4 and 40 µM) on (**C**) CA_S16E_ multimerization in higher sodium concentration (the ratio of 150 mM to 5M sodium is four to six), and (**D**) CA_T19A_ multimerization (the ratio of 150 mM to 5M sodium is five to five) are shown. Representative data is shown from three independent experiments.

### ACAi-028 potentially interacts with the binding pockets of CA in the hexameric state

ACAi-028 exerted anti-HIV-1 activity in the early stages of the HIV-1 life cycle. In the early stage, CA proteins constitute a capsid lattice (HIV core), which is composed of approximately 250 hexamers and exactly 12 pentamers (41). To confirm the location and space of the binding pocket in a CA hexamer, we analyzed a CA hexamer model in complex with, or without, ACAi-028, based on CA crystal structures ([PDB] accession number 3H4E (42), and 5MCX (32)). In the model, the ACAi-028 can bind to the pocket located at inside the CA hexamer ([PDB], 3H4E) (Fig. 8A). Additionally, the ACAi-028 binding profile to the pocket in a CA dimer extract from the CA hexameric state ([PDB], 5MCX) was predicted using our docking model. ACAi-028 can putatively interact with the pocket in the CA dimer of the CA hexameric state, without affecting the paired CA monomer, via two H-bonds with L43 and E45 (Fig. 8B). This suggests that ACAi-028 can potentially interact with the binding pockets of the CA hexameric state, in the HIV-1 core. Taken together, ACAi-028 is a novel capsid inhibitor that binds to the new hydrophobic pocket in CA-NTD, thereby inhibiting the early stage of HIV-1 replication.

**Figure 8.**
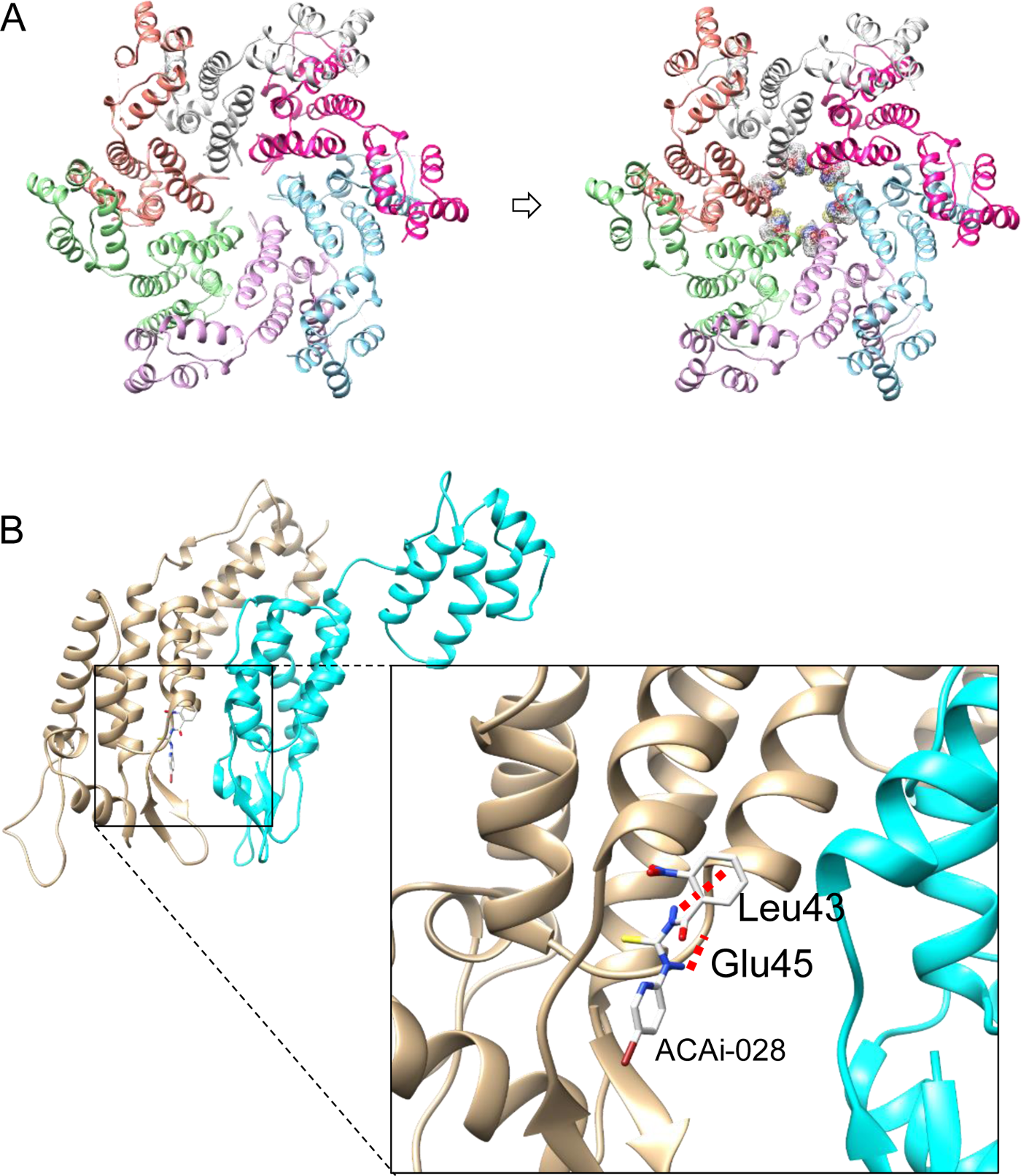
Interaction of ACAi-028 with the binding pockets of CA in the hexameric state. **(A)** Left panel shows the structure of a CA hexamer ([PDB] accession number 3H4E). CA dimer extracted from the CA hexamer was used in the flexible docking simulation. Right panel shows the CA hexamer with docking poses of ACAi-028. **(B)** The structure of a CA dimer extracted from a CA hexamer ([PDB] accession number 5MCX) and the docking result of ACAi-028 to the CA dimer is shown. ACAi-028 formed two H-bond interactions with the main-chains of Leu43 and Glu45 residues which are located at the monomer-monomer interface of one CA monomer in the CA dimer extracted from the CA hexamer. Docking simulations were performed with SeeSAR and FlexX version 10. Molecular graphics was performed with UCSF Chimera.

## Discussion

Based on the results described above, we concluded that ACAi-028 is a small molecular CA inhibitor of HIV-1 that interacts with CA via the novel region, which has not been previously reported (Fig. 9 and Table 4). The ACAi-028 binding pocket is formed by key residues, namely at Gln13, Ser16, and Thr19, which constitute the β-hairpin end, flexible linker, and front edge of α-helix 1 (Fig.1). To understand the mechanisms of action of CA inhibitors previously described (43), we categorized several candidate CA inhibitors based on their molecular characterization into groups A, B, C, and D (Fig. 9 and Table 4). CAP-1 (group A) is a late-stage inhibitor, without early stage inhibition of the HIV-1 life cycle (13, 14), possibly because the hexameric association of CA proteins in the mature HIV-1 core requires tight molecular packing, prevent access of CAP-1 to this pocket. Indeed, the other small molecules, including BD-1 and BM-1 (18), which share the CAP-1 binding region, are all late-stage inhibitors. Therefore, the inhibitory mechanism of group A is likely different from that of ACAi-028. PF74 (group B) interacts with a region distinct from ACAi-028 and CAP-1, as shown in Fig. 9 and Table 4, and binds to the pocket formed between CA-NTD and the CA-CTD of an adjacent CA monomer. PF74 is known to inhibit both the early and late stages of the HIV-1 life cycle (15), whereas ACAi-028 only inhibited the early stage (Fig. 2 and 3). As shown in the CA multimerization and DSF assay (Fig. 5), ACAi-028 has a significant effect on the CA multimerization and thermal stability, which are completely opposite to PF74, suggesting that ACAi-028 and PF74 represent different classes of CA inhibitors (Fig. 9 and Table 4). I-XW-053 (19) (group C) binds to, and disrupts CA NTD-NTD interactions in CA hexamers during the early stages of HIV-1 infection. By SPR analysis of I-XW-053 binding with CA mutants, the proposed binding sites of I-XW-053 were found to involve I37 and R173, which the amino acids identified within the target binding region of ACAi-028. Group C includes C1 (20), a late-stage assembly inhibitor that interacts with the CA-NTD residues at E98, H120, and I124 residues. Ebselen (group D) is a covalent inhibitor of HIV-1 CA by forming a selenosulphide bond with C198 and C218 residues in the CA-CTD. ACAi-028 possesses distinct properties, in comparison to the inhibitors of groups C and D, suggesting that ACAi-028 targeting the hydrophobic pocket is likely to be different from any previously discovered CA inhibitors.

**Figure 9.**
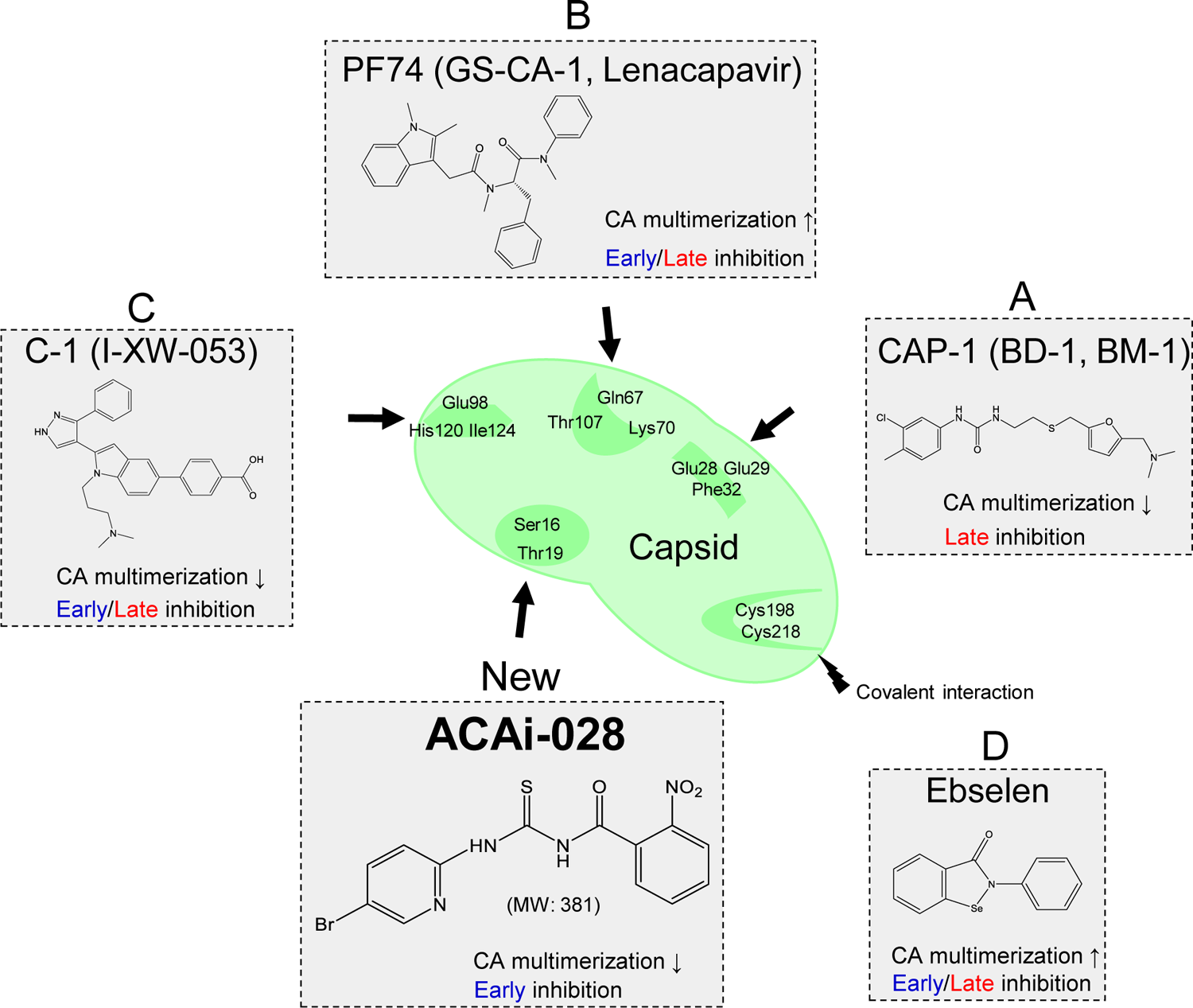
Profiles of ACAi-028 and CA inhibitors. ACAi-028 and representative CA inhibitors previously reported are categorized into groups new, A, B, C, and D. Names, chemical structures, effects of CA multimerization, inhibition stages of the HIV-1 life cycle, putative and representative binding regions of the CA inhibitors are shown.

**Table 4:** Comparison of ACAi-028 binding position and mechanism with other CA inhibitors.

| CA | Group | Name | Binding region | C.M. | Stage of inhibition | Ref. |
| --- | --- | --- | --- | --- | --- | --- |
| NTD | New | ACAi-028 | Q13, S16, T19 | ↓ | Early |  |
|  | A | CAP-1 | E28, E29, F32, V59, H62 | ↓ | Late | 13, 14 |
|  |  | BD-1, BM-1 | F32, H62 | ↓ | Late | 20 |
|  | B | PF74 | Q67, K70, T107 | ↑ | Early/Late | 15 |
|  |  | (GS-CA1)<br>Lenacapavir | Q67, K70, T107 | ↑ | Early/Late | 17, 18 |
|  | C | I-XW-053 | I37 | ↓ | Early | 21 |
|  |  | C1 | E98, H120, I124 | ↓ | Late | 22, 44 |
| CTD | D | Ebselen | C198, C218 | ↑ | Early/Late | 16 |
|  | B | PF74 | Y169, L172, R173,Q179 | ↑ | Early/Late | 15 |
|  |  | (GS-CA1)<br>Lenacapavir | Y169, L172, R173,Q179 | ↑ | Early/Late | 17, 18 |
|  | C | I-XW-053 | R173 | ↓ | Early | 21 |

**C.M.**, CA Multimerization ↑ Increase, ↓ Decrease **Ref.**, Reference.

Amino acid substitutions that alter CA multimerization have been previously reported (38). Substitution of amino acid residues at S16 and T19 altered the CA multimerization; S16E mutants decreased the CA multimerization, whereas T19A mutants increased the CA multimerization. Additionally, E45A substitution, E45 residue is a residue predicted to be located in theACAi-028-binding pocket, is known to increase the CA multimerization (45). These amino acids might constitute an important site for CA dimerization in the formation of CA hexamers (Fig. 8). The binding of ACAi-028 to the pocket could potentially interfere with CA dimerization, because this pocket is located within an inward facing portion of the CA hexamer (Fig. 8A), which is supported by the reduction of CA multimerization in the presence of ACAi-028 (Fig. 5A). To investigate the binding profile of ACAi-028 to CA-NTD, we produced crystals of CA-NTD_1-146/Δ87-99G_ proteins, according to a previous report (15), and utilized this structure for the preparation of a docking model for ACAi-028 binding to CA. Unfortunately, when crystals of CA-NTD_1-146/Δ87-99G_ were produced in the presence of ACAi-028, significant precipitations occurred without the emergency of a crystal in the droplet, suggesting that ACAi-028 may cause CA proteins instability or destabilization of the CA protein. Thus, we were unable to acquire actual binding profiles of ACAi-028 to CA. Moreover, when we have selected HIV-1 variants resistant to ACAi-028, with increasing concentrations of up to 20 µM, we did not detect ACAi-028-related strong resistant mutations in the CA (Gag) regions, suggesting that ACAi-028 might still possess an unknown mechanism for HIV-1 inhibition. This represents a limitation of the present study, and requires further investigation.

Recently, lenacapavir was reported to be a powerful anti-HIV compound, with broad-spectrum inhibition, even against multidrug-resistant HIV-1, HIV-2, and SIV (21), for long-acting HIV-1 treatment (21, 22) (Phase 2/3 CAPELLA trial). Furthermore, lenacapavir has also been predicted to be effective in HIV prevention as HIV pre-exposure prophylaxis (PrEP). These findings strongly suggest that CA is an attractive therapeutic target for the development of novel antivirals.

In conclusion, we have identified ACAi-028 as a small molecular anti-HIV-1 CA inhibitor that targets a novel hydrophobic CA-NTD pocket, exerting the early-stage inhibition of the HIV-1 life cycle with EC_50_ of 0.55 µM. Further research is underway to understand the role of this region in HIV-1 replication. The novel hydrophobic pocket identified here, should be a viable target for the development of new synthetic CA inhibitors. Furthermore, ACAi-028, a potent CA inhibitor targeting this novel pocket, could have valuable therapeutic and research applications.

## Materials and Methods

### Cells and Viruses

MT-2 and MT-4 cells (Japanese Collection of Research Bioresources Cell Bank; JCRB Cell Bank, Japan) were cultured in RPMI1640 medium (Gibco, Thermo Fisher Scientific, USA) with Fetal Bovine Serum (FBS, Gibco, Thermo Fisher Scientific, USA), Penicillin (P), and Kanamycin (K). 293T, Li 7, HLE, and COS7 cells obtained from JCRB Cell bank and TZM-bl cells obtained from the NIH AIDS Research and Reference Reagent Program were cultured in low glucose DMEM with L-Glutamine and Phenol Red (Fujifilm Wako Pure Chemical Corporation, Japan), as well as FBS, P, and K. PHA-PBMCs were derived from a single donor in each independent experiment. The research protocols described in the present study were carried out in accordance with relevant guidelines and regulations, and were approved in Ethics Committee for Epidemiological and General Research at the Faculty of Life Sciences, Kumamoto University. The HIV-1 strains used in our experiments have already been established previously, including HIV-1_NL4-3_, HIV-1_LAI_, HIV-1_Ba-L_, HIV-1_JR-FL_, HIV-1_ERS104pre_ which was isolated from clinical HIV-1 strains of drug-naive patients with AIDS (46), and HIV-1_MDR/B_ which was originally isolated from AIDS patient who had received 9 anti-HIV-1 drugs over the 34 months and was genotypically and phenotypically characterized as multi-drug-resistant HIV-1 variant (47, 48). HIV-1_VSV-G dENV_ was produced by co-transfection of pCMV-VSV-G vector (addgene) and pNL_4-3 dENV_ with deleted Kpn1-Nhe1 site in the Env region into 293T cells.

### Plasmid constructs

Full-length CA sequences derived from pNL_4-3_ was introduced to pET30a vectors (Novagen-Merck KGaA, Germany), producing a pET30a CA vector. The site-directed mutagenesis was performed using PrimeSTAR**^®^** Max (Takara Bio Inc. Japan) to introduce S16E, T19A, E45A, and M185A into the pET30a CA, producing pET30a CA_S16E_, CA_T19A_, and CA_M185A_ vectors, respectively. CA-NTD_146_ deletion sequence carrying a single glycine residue instead of CypA-BL (residues 87–99) (the amino acids sequence is based on (15)) was introduced to pET30a vectors 6xHis at N-terminus, producing a pET30a His-CA _1-146/Δ87-99G_ vector

### Protein expression and purification

CA proteins were produced from the pET30aCA in E. coli Rosetta^TM^ (DE3) pLysS Competent Cells (Novagen) grown in LB medium supplemented with Kanamycin and chloramphenicol at 37°C, and induced with 1.0 mM Isopropyl β-D-1-thiogalactopyranoside (IPTG) for 3-5 hr at 37°C. The bacterial cells were harvested and stored at −80°C. The pellets of CA were resuspended and sonicated in CA buffer (50 mM Tris-HCl pH 7.5, 150 mM NaCl, 5 mM 2-mercaptoethanol (BME)) supplemented with 0.5 mM phenylmethylsulfonylfluoride. The lysates were cleared by centrifugation for 15 min at 3,500 rpm at 4°C. After 5M NaCl was added to the supernatants, the samples were cleared by centrifugation for 15 min at 3,500 rpm again. The precipitate was resuspended in the CA buffer. The sample was precleared for 15 min at 15,000 rpm at 4°C, and the supernatants were filtered through a 0.45 μm filter.

The CA proteins were loaded onto a HiLoad 16/60 Superdex 200 column (GE Healthcare, USA), and were eluted with CA loading buffer (20 mM Tris-HCl pH 7.5, 150 mM NaCl, 5 mM BME) using AKTAprime plus (GE Healthcare). CA monomer fractions were concentrated using Amicon Ultra-10K device (Merck Millipore) in the CA buffer. The proteins concentration was determined using a BCA Protein Assay Reagent Kit (Thermo Fisher Scientific) and stored at −80°C.

### *In silico* simulation and docking model

The crystal structures of CA-NTD_1-146/Δ87-99G_ produced by our method and various full-length CA proteins ([PDB] accession number 4XFX, 3H4E, and 5MCX) from the RCSB Protein Data Bank (http://www.rcsb.org) are utilized for the docking simulation. Hydrogens were added to 2D structure of ACAi-028, and the structures were energy-minimized with the MMFF94x force field as implemented in MOE (Chemical Computing Group, Quebec, Canada). All docking simulations were performed with SeeSAR and FlexX version 10 (BioSolveIT GmbH, Sankt Augustin, Germany). Molecular graphics and analysis were performed with UCSF Chimera (https://www.rbvi.ucsf.edu/chimera).

### Fusion assay

Fusion assay was performed previously (31). In brief, 293T cells were transfected with pHIV-1_NL4-3_ Tat with or without pHIV-1_NL4-3_ Env while COS-7 cells were transfected with CD4, CXCR4, and LTR-Luciferase. After 24 hr of incubation at 37°C and 5% CO_2_, the transfected 293T cells were mixed with the transfected COS-7 cells in the presence or absence of tested compounds for 6 hr. The luciferase of the samples was detected using Firefly Luciferase Reporter Assay Kit I (PromoCell GmbH, Germany) and the luciferase intensity was normalized to the negative control. Ratios of luciferase intensity of the samples were compared.

### Time-of-addition assay

Time-of addition assay is previously reported (32). Briefly, TZM-bl cells (5 × 10^4^/mL) were subjected to HIV-1_LAI_ (50 ng/mL of p24) in 96-well white plates. Drugs at the indicated concentrations were also added to the set of wells demarcated for 0 hr. After 2 hr of incubation at 37°C and 5% CO_2_, The supernatant was removed and the cells were washed once to remove all traces of the virus. Drugs at the respective concentration were added back to the cells marked for 0 hr as well as the ones marked for 2 hr. Subsequently, every 2 hr after, drugs at the correct concentration was added to the next each well until 10 h after incubation. The cells were incubated at 37°C and 5% CO_2_ until 48 hr. Luciferase intensity was measured using FluoSTAR Omega (BMG labtech GmbH, Germany).

### TZM-bl assay

TZM-bl cells were obtained from the NIH AIDS Research and Reference Reagent Program. TZM-bl assay was performed in 96-well white plate using the Firefly Luciferase Reporter Assay Kit I. Briefly, the supernatant medium was removed and lysis buffer added to the samples. After the samples were shaken for 25 minutes, D-luciferin was added to each well. After shaking, the luciferase intensity was measured using FluoSTAR Omega.

### Virus quantification

Virus samples were measured by an HIV-1 p24 enzyme-linked immunosorbent assay (ELISA) using Lumipulse G1200 (Fujirebio Inc, Japan), and normalized to determine the viral concentration.

### Reverse Transcriptase Assay

Colorimetric reverse transcriptase assay was performed (Reverse Transcriptase Assay colorimetric, Roche, Switzerland). Briefly, recombinant HIV-1 reverse transcriptase was added to tested compounds dissolved in the lysis buffer and subsequently incubated at 37°C and 5% CO_2_ for 3-4 hr. After washing, peroxidase-conjugated anti-digoxigenin antibody solution were added to the samples and incubated for 1 hr at 37°C and 5% CO_2_. After washing, peroxidase substrate ABTS solution with enhancer was added to the samples. The optical density was measured at 405 nm using a Versamax microplate reader (Molecular Devices, USA).

### Western blotting

Western blotting was described previously (40). Briefly 293T cells were plated and incubated for 24 hr at 37◦C in 5% CO_2_. Cells were transfected with pNL_4-3_ vectors using Attractene Transfection Reagent (QIAGEN, Germany). After 8 hr, the medium was changed and the tested compounds were added, and then incubated for 48 hr. Subsequently, the viruses were filtered, purified by ultracentrifugation in 15% sucrose–phosphate-buffered saline (PBS), normalized by the p24 levels, and stored in PBS at −80°C. The cells were lysed in M-per buffer (Thermo Fisher Scientific) supplemented with Halt Protease Inhibitor Cocktail (Thermo Fisher Scientific). The samples were titrated using BCA Protein Assay Kit and stored at −80°C. The samples were prepared and separated with SDS-PAGE (5-20% Extra PAGE One Precast Gel; nacalai tesque) and transferred onto a nitrocellulose membrane. The samples were detected with anti-HIV-1 Gag (p55 + p24 + p17) antibody (catalog number ab63917; Abcam), anti-HIV-1 IN antibody (catalog number ab66645; Abcam), second mouse or rabbit antibody (MBL co., LTD.), and anti-beta actin antibody (HRP conjugated) (Abcam), and then visualized using SuperSignal WestPico Chemiluminescent Substrates (Thermo Fisher Scientific).

### CA multimerization assay

CA multimerisation assays were performed as previously reported (38). Compound was added to 30 µM of CA protein in 50 mM phosphate buffer at pH 8.0 supplemented with 50mM NaCl. Capsid assembly was initiated by addition of 50 mM sodium phosphate at pH 8.0 supplemented with 5M NaCl. Optical density at 350 nm was measured using FLUOstar Omega (BMG labtech) for 1 hr.

### DSF

DSF method was previously described (39, 40). In brief, recombinant CA proteins (25 µM) were prepared in PBS. After CA treatment with tested compounds for 5-8 hr on RT, SYPRO Orange (Life Technologies) was added to the samples (final concentration of SYPRO orange: 5×). The samples were successively heated from 25 to 95°C, and the increasing fluorescence intensities were measured by the real-time PCR system 7500 Fast (Applied Biosystems, Thermo Fisher Scientific). Data indicated is as a relative ratio between minimum and maximum intensity of SYPRO orange from 25 to 95°C detected for each sample.

### ESI-MS

ESI-MS protocol was previously described (37). In brief, MS spectra of CA in the presence of ACAi-028 were obtained using an electrospray ionization (ESI) quadrupole time-of-flight (QTOF) mass spectrometer (impact II, Bruker Daltonics). Each sample solution in native condition was introduced to the ESI-QTOF mass spectrometer through an infusion pump at a flow rate of 3.3 μL/min. To detect the denatured samples, analysis was done using the QTOF mass spectrometer equipped with a Captive Spray electrospray ionization platform with liquid chromatography (Ultimate 3000 HPLC, Thermo Fisher scientific). The following ion source parameters have been applied: dry Heater: 150°C, dry Gas: 8.0 L/min, capillary voltage: 1000 V, End plate offset: −500 V. MS scans have been acquired at a spectra rate of 1 Hz at a mass range from 100 to 3000 m/z. Molecular weights by protein deconvolution were determined using Data Analysis 4.4 (Bruker Daltonics). M.W. of CA proteins was calculated using Peptide Mass Calculator v3.2 (http://rna.rega.kuleuven.be/masspec/pepcalc.htm)

### Crystallization and X-ray data collection

The crystallization procedure was performed according to methods in a previous report (15) ([PDB] 2XDE). Briefly, crystallization was performed by the hanging-drop vapor diffusion method using EasyXtal 15-well tools (QIAGEN). CA-NTD _1-146/Δ87-99G_ proteins were expressed and purified as described above. The protein concentration was adjusted to 2 mg/ml. The reservoir solution consists of 100 mM phosphate-citrate pH 4.2, 200 mM NaCl, and 20% PEG 8000. The crystals reached 0.2 to 0.4 mm within 1 week at 10°C. The crystals were transferred to a reservoir solution supplemented with 25% glycerol and flash-frozen at 100 K. Then, X-ray diffraction experiments were carried out. Data collection and refinement statistics were subsequently examined.

### Drug Susceptibility Assay

The susceptibility of HIV-1_LAI_ and HIV-2_ROD_ to ACAi-028 and control drugs/compounds were determined as previously described (49). Briefly, MT-2 cells (10^4^/ml) were exposed to 100 × 50 % tissue culture infectious dose (TCID_50_) of HIV-1_LAI_ or HIV-2_ROD_ in the presence or absence of various concentrations of compounds in 96-well plates and were incubated at 37°C for 7 days. After incubation, 100 µl of the medium was removed from each well, 3-(4,5-dimetylthiazol-2-yl)-2,5-diphenyltetrazolium bromide (MTT) solution was added to each well in the plate, followed by incubation at 37°C for 1.5-4 hr. After incubation to dissolve the formazan crystals, acidified isopropanol containing 4% (v/v) Triton X-100 was added to each well and the optical density measured in a kinetic microplate reader (Vmax; Molecular Devices, Sunnyvale, CA). All assays were performed in duplicate. 50% cytotoxic concentration (CC_50_) of compounds/drugs for each cell line was also evaluated by MTT assay. To determine the sensitivity of primary HIV-1 isolates to compounds, PHA-PBMC (10^6^/ml) were exposed to 50 TCID_50_ of each primary HIV-1 isolate and cultured in the presence or absence of various concentrations of drugs in 10-fold serial dilutions in 96-well plates. In determining the drug susceptibility of certain laboratory HIV-1 strains, MT-4 cells were employed as target cells as previously described (50, 51) with minor modifications. In brief, MT-4 cells (10^5^/ml) were exposed to 100 TCID_50_ of drug-resistant HIV-1 strains in the presence or absence of various concentrations of compounds and were incubated at 37°C. On day 7 of culture, the supernatants were harvested and the amounts of p24 (CA) protein were determined by using Lumipulse G1200. Drug concentrations that suppressed the production of p24 Gag protein by 50% (EC_50_; 50% effective concentration) were determined by comparison with the p24 production level in drug-free control cell culture. PHA-PBMCs were derived from a single donor in each independent experiment.

### Compounds

ACAi-028 was purchased from ChemBridge (San Diego, CA, USA). AZT, PF74, and EFV were purchased from Sigma-Aldrich, Ebselen from AdipoGen Life Sciences, and AMD3100 from Selleck Chemicals. Atazanavir (ATV) was kindly provided by Bristol Myers Squibb (New York, NY).

## Acknowledgements

We thank Teruya Nakamura for crystal determination and Sachiko Otsu for technical assistance of the experiments. We also thank the beamline staff at Photon Factory and SPring-8 for their help in the data collection.

This work was supported by Japan Society for the Promotion of Science KAKENHI Grant Numbers JP18K08436 (HN) and JP15K09574 (MA). Development of Novel Drugs for Treating HIV-1 Infection and AIDS, and the grants from the Sumitomo Electric Group CSR Foundation (MA).

TN and MA designed and TC, TN, NT, and MA performed all the experiments. NT and MM discussed the data and supported preparation of the MS. MA and HN supervised, managed the project, and acquired the necessary funding. TC, TN, and MA wrote and MM and HN edited the MS. All authors read, commented on, and approved the final manuscript.

We declare that we have no conflicts of interest.

